# Oncogenic *KRAS* Mutations Confer a Unique Mechanotransduction Response to Peristalsis in Colorectal Cancer Cells

**DOI:** 10.1101/2024.05.07.593070

**Authors:** Abigail J. Clevenger, Claudia A. Collier, John Paul M. Gorley, Maygan K. McFarlin, Spencer C. Solberg, E. Scott Kopetz, Amber N. Stratman, Shreya A. Raghavan

**Author notes:** Address Correspondence to: Shreya A. Raghavan, PhD (she/her) 5016 Emerging Technologies Building 3120 TAMU College Station TX 77843.

## Abstract

Colorectal cancer (CRC) tumors start as precancerous polyps on the inner lining of the colon or rectum, where they are exposed to the mechanics of colonic peristalsis. Our previous work leveraged a custom-built peristalsis bioreactor to demonstrate that colonic peristalsis led to cancer stem cell enrichment in colorectal cancer cells. However, this malignant mechanotransductive response was confined to select CRC lines that harbored an oncogenic mutation in the *KRAS* gene. In this work, therefore, we explored the involvement of activating *KRAS* mutations on peristalsis-associated mechanotransduction in CRC. Peristalsis enriched the cancer stem cell marker LGR5 in *KRAS* mutant (G13D, etc.) lines, in a Wnt-independent manner. Conversely, LGR5 enrichment in wild type *KRAS* lines exposed to peristalsis were minimal. LGR5 enrichment downstream of peristalsis translated to increased tumorigenicity *in vivo* in *KRAS* mutant vs. wild type lines. Differences in mechanotransduction response was additionally apparent via unbiased gene set enrichment analysis, where many unique pathways were enriched in wild type vs. mutant lines, in response to peristalsis. Interestingly, peristalsis also triggered β-catenin nuclear localization independent of Wnt, particularly in *KRAS* mutant lines. The central involvement of KRAS in the mechanotransductive responses was validated via gain and loss of function strategies. β-catenin activation and LGR5 enrichment downstream of peristalsis converged to the activation of the MEK/ERK kinase cascade, that remains active in cells that harbor oncogenic *KRAS* mutations. Taken together, our results demonstrated that oncogenic *KRAS* mutations conferred a unique peristalsis-associated mechanotransduction response to colorectal cancer cells, resulting in cancer stem cell enrichment and increased tumorigenicity. These mechanosensory connections can be leveraged in improving the sensitivity of emerging therapies that target oncogenic KRAS.

## Introduction

Colorectal cancer (CRC) is the second most common cause of cancer related deaths in the United States^1, 2^. To best improve patient outcomes with targeted therapeutics, it is important to first understand the initial stages of CRC development. CRC tumors begin as a precancerous polyp on the inner lining of the colon or rectum^3, 4^. Precancerous polyps can switch to progressive and malignant carcinomas (CRC). Our knowledge of why some polyps become malignant CRC is still growing. In CRC, mutations in the Kirsten rat sarcoma virus (*KRAS*) gene typically occur in the early stages of the polyp to malignant carcinoma transition^5–8^. Importantly, clinical pathology of CRC tumors indicates that 35-40% of CRC patients harbor a *KRAS* mutation, making it an appealing target to improve patient prognosis^9–11^.

In homeostasis, the *KRAS* gene relays signals from upstream growth factor receptors to downstream effector pathways such as RAF/MEK/ERK, MAPK, and PI3K, among others. These pathways control important processes such as cell cycle progression, cell survival, cell polarity and movement, actin cytoskeletal organization, and extracellular signal transduction^12, 13^. Oncogenic activating mutations, such as G13D, G12D, and G12C, in the *KRAS* oncogene drive cell transformation and uncontrolled cell division^14, 15^ by locking *KRAS* in an active GTP-bound conformation, thereby driving constant downstream pathway activation^16, 17^. Clinical analysis of CRC tumor samples indicates that activating *KRAS* mutations are associated with increased metastasis, CRC recurrence, and overall poorer patient survival^6, 18–20^.

Oncogenic mutations in the *KRAS* gene are known to amplify cancer cell mechanosensing in breast and pancreatic cancers, particularly as it relates to transmission of stiffness from the extracellular matrix^21^. *KRAS* driven changes in mechanosensing were directly related to altered cytoskeletal tensional homeostasis in cancer cells. Interestingly, in CRC patient samples, *KRAS* mutated tumors were stiffer than non-mutated tumors^22^. This led us to hypothesize that CRC cells harboring activating oncogenic *KRAS* mutations will have uniquely altered mechanotransduction leading to malignant CRC progression.

In the CRC tumor microenvironment, cells in the intestinal lumen, including cells in precancerous polyps, are continuously exposed to mechanical forces associated with colonic peristalsis^23, 24^. In previous studies, we developed and validated a peristalsis bioreactor that mimics the forces associated with colonic peristalsis^25^. Using the peristalsis bioreactor, we demonstrated that HCT116 *KRAS^G13D^* cells exposed to peristalsis increased cancer stem cell markers relative to static culture^26^. HCT116 cells also developed invasive signatures downstream of peristalsis, indicating malignant progression. Motivated by our previous studies and the amplified mechanosensing reported in other tumors harboring *KRAS* mutations, we hypothesized that activating *KRAS* mutations will alter peristalsis-associated mechanotransduction in CRC.

Leveraging the peristalsis bioreactor, multiple cell lines and patient-derived xenograft lines were employed to test the hypothesis that activating *KRAS* mutations change mechanotransduction via cancer stem cell enrichment. Our focus was on the Leucine rich repeat containing G protein couple receptor, LGR5, a cancer stem cell marker which our previous work strongly demonstrated was downstream of Wnt activation^26^. Here, we characterize changes in the Wnt pathway as a function of *KRAS* mutation status in response to peristalsis. To confirm the involvement of mutant *KRAS* in the altered mechanotransduction response to peristalsis, we employed gain and loss of function techniques for *KRAS* and its downstream effectors. This work identifies a connection between activating *KRAS* mutations and altered cellular response to peristalsis. Results from this work will be vital to improving our understanding of how activating *KRAS* mutations impact CRC malignant progression via mechanotransduction.

## 2. Methods

### 2.1 Materials

Cell culture reagents were purchased from ThermoFisher Scientific (Waltham, MA) unless otherwise specified. Cell lines were purchased from American Type Culture Collection (ATCC; Manassas, VA) unless otherwise specified. Polydimethylsiloxane (PDMS) was purchased from DOW Chemical (Midland, MI). All other chemical reagents were purchased from Sigma Aldrich (St. Louis, MO), unless otherwise indicated. Antibodies used for cellular staining were purchased from Santa Cruz Biotechnology (Dallas, TX), unless otherwise indicated. Custom-made oligos, including CRISPR reagents, were purchased from Integrated DNA Technologies (Coralville, IA). All other molecular biology-grade reagents were purchased from ThermoFisher Scientific (Waltham, MA).

### 2.2 Cell Culture

Three characterized colorectal cancer cell lines HCT116, LS174T, and RKO were used in this work. Additionally, three patient derived xenograft lines, PDX1, PDX2, and PDX3, established from primary patient samples^27^, were utilized to assess the effects of peristalsis (PDX lines were gifts from the Kopetz lab at the University of Texas MD Anderson Cancer Center). Dulbecco’s Modified Eagle Medium (DMEM), Eagle’s Minimum Essential Media (EMEM, ATCC), and Roswell Park Memorial Institute (RPMI) 1640 Medium supplemented with 10% heat-inactivated fetal bovine serum (Peak Serum, Inc., Wellington, CO) and 1X Antibiotic-Antimycotic solution were used as the primary growth media for HCT116, LS174T and RKO, and PDX cells, respectively. All cells were cultured in standard 2D tissue culture flasks and treated with 0.25% trypsin to dissociate adherent cells. Molecular characterization of the cell and PDX lines are presented in **Table 1**.

**Table 1:**
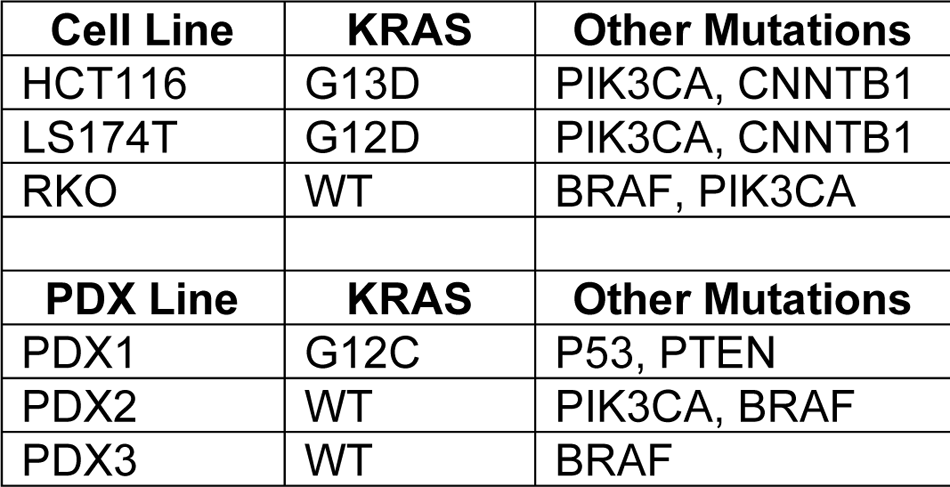
Mutation status of all cell lines and PDX lines used in this work (WT = Wild Type)^28, 29^.

### 2.3 Preparation of Cell Seeded Membranes and Bioreactor Assembly

Polydimethylsiloxane (PDMS) membranes were prepared at a 10:1 ratio using previously established protocols^25, 26^. To maximize seeding of cells on PDMS, the cell seeding area of the membranes were coated with 200 µg/mL of Collagen I. Cells were seeded onto PDMS at monolayer confluency of 500,000 cells/mL and allowed to adhere at 37°C for 4 hours. Cell attachment was confirmed via visual inspection with a cell culture microscope.

Operation of the peristalsis bioreactor was followed as previously reported^25, 26^. Briefly, media was removed from each cell-seeded PDMS membrane. Then, cell-seeded peristalsis membranes were placed into the peristalsis bioreactor bottom while static membranes were maintained in cell culture dishes with 1 mL of fresh media. The bioreactor was assembled using commercially available zip ties and connected to the peristalsis pump. The assembled set up was placed into the 37°C incubator and connected to an Arduino that ran a pre-programmed code with the following parameters: 0.4 Pa shear and 15% cyclic strain at 12 rpm^25, 26^. Cells were incubated, either as static controls or stimulated in the peristalsis bioreactor at 37°C for 24 hours and then collected for downstream analysis.

Control untreated conditions were maintained in static membranes, or peristalsis bioreactors in their respective growth medium. To determine the effects of inhibiting Wnt in some experimental conditions, a porcupine-selective pan-Wnt inhibitor (LGK974, 1 µM; Med Chem Express, Monmouth Junction, NJ) was incorporated into the growth media. To study the effect of MEK inhibition, after cells adhered to PDMS for 4 hours, media was removed from cell-seeded PDMS and growth media supplemented with Selumetinib (50 nM; Selleck Chemicals, Houston, TX) was added. Following overnight incubation, Selumetinib (50 nM) was incorporated into the growth media for appropriate peristalsis conditions. A list of experimental media variations is provided in **Table 2**.

**Table 2:** Experimental media formulations.

| Experimental Condition | Additional Media Components |
| --- | --- |
| Control | None |
| Wnt inhibition | LGK974 1 µM |
| MEK inhibition | Selumetinib 50 nM |

### 2.4 Flow Cytometry Analysis of LGR5 Cancer Stem Cell Phenotype

Following 24-hour maintenance in static or exposure to peristalsis conditions, cells were detached using 0.25% trypsin and collected from PDMS into single cell suspensions in FACS buffer (Phosphate Buffered Saline (PBS) supplemented with 2% fetal bovine serum). Flow cytometry methods were performed using protocols optimized previously for cancer cells^26, 30, 31^. Briefly, cells were incubated with AlexaFluor488-LGR5 antibody (R&D Systems, Minneapolis, MN), or an isotype matched AlexaFluor-488 antibody (R&D Systems) for 30 minutes at 37°C. Unbound antibody was removed by washing and resuspending in fresh FACS buffer and cells were analyzed on the Attune NxT flow cytometer (ThermoFisher Scientific). Isotype controls were used to establish a gating strategy by cutting off a background gate at 0.5% (gating strategy is demonstrated in **Supp.** Fig. 1). Based on the background gate, the percentage of cells expressing LGR5 was determined. Comparisons were drawn between static and peristalsis conditions.

### 2.5 Sample Collection and Processing of RNA-Seq Data

Cells were exposed to static or peristalsis conditions for 24 hours; RNA was extracted directly from the bioreactor membranes following manufacturer’s instructions from the RNeasy Mini Kit (Qiagen, Hilden, Germany). RNA concentration was assessed using the Qubit Fluorometer (Life Technologies; Carlsbad, CA) and RNA quality was evaluating used the Tape Station (Agilent; Santa Clara, CA). Total RNA samples with adequate quality (RIN ≥6.0) and amount (≥5 ng/µl with >500ng) were sent to Azenta Life Sciences (Burlington, MA) for RNA-Sequencing.

Raw data files in FASTQ format were generated from the Illumina sequencer obtained from Azenta Life Sciences. Initial preprocessing involved quality control using FastQC software to trim adapters and low-quality bases. Clean reads were then mapped to the Human genes GRCh38.p13 reference genome available on ENSEMBL using the STAR aligner v.2.5.2b. Differential expression analysis was conducted using DESeq2 (R package), accounting for biological replicates, to identify genes with a fold change greater than 2 and an adjusted p-value (FDR) less than 0.1.

#### 2.5.1 Weighted Gene Correlation Network Analysis (WGCNA)

K-means clustering was performed to visualize the expressed genes using iDEP tool suit^32^. Functional enrichment analysis utilized EnRichGO (R library), on select genes grouped through WGCNA with minimum module size of 20 and a soft threshold of 15 to best fit the network structure using pickSoftThreshold function, considering the gene’s association with biological processes. All statistical analyses were conducted in R studio (v2023.12.1+402) and iDEP with significance thresholds adjusted based on the study design and corrected for multiple testing where applicable.

#### 2.5.2 Gene Set Enrichment Analysis (GSEA)

Normalized count files from DESeq2 analysis performed by Azenta Life Sciences were populated into the Gene Set Enrichment Analysis Software (https://www.gsea-msigdb.org/gsea/downloads.jsp)^33, 34^. Analysis was run with two different gene sets: i) Gene Ontology Biological Process (GOBP)^35, 36^ and ii) Hallmark^37^ MSigDB gene sets. For each analysis, gene set files, gene count files, and phenotype label files were loaded in GSEA software^38^. To reach a normalized enrichment score (NES), gene set permutations were conducted 1000 times^39^. The minimum and maximum criteria for gene sets selection were 1 and 1000 genes, respectively. The NES normalizes the enrichment score to the size of the gene set. To account for false positives in multiple testing, an NES was considered significant with a false discovery rate (FDR) less than 0.25^38^. A higher NES indicates more positively enriched genes in the gene set.

### 2.6 Assessment of Tumorigenicity in NSG Mice

HCT116 *KRAS^G13D^* and RKO *KRAS^WT^* cells were either maintained as static controls or stimulated with peristalsis in the bioreactor for 24 hours. Cells were harvested using 0.25% trypsin and prepped for subcutaneous injection into NOD.Cg-Prkdc^scid^ Il2rg^tm1Wjl^/SzJ (NSG) mice. These methods were adapted from previous protocols established by Raghavan, et al.^31^. Animal experiments were performed following approval by the Texas A&M University Institutional Animal Care and Use Committee under protocol 20204-0040 D. Three male mice and three female mice were used per test condition (randomly assigned to static and peristalsis groups). Following harvest, a total cell count was obtained and then adjusted to a concentration of 140,000 cells per 50 µL of media. Ice cold Matrigel was added to cell concentrations at a ratio of 1:1 Matrigel:media. Each mouse received two subcutaneous flank injections (one on either side of the body) of 100 uL each (i.e. 140,000 cells per injection).

For two weeks following injections, mouse weight was monitored twice a week and injection sites were palpated for any tumor presence. Once tumors were present, measurements were obtained three times a week using a caliper to record tumor volumes (**Eq. 1**) and mouse weight was continually monitored.

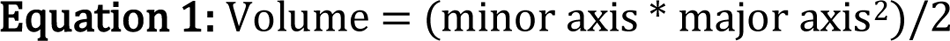

Tumor volumes were recorded until tumors reached an end point of 2,000 to 2,500 mm^3^ and then mice were euthanized. Tumors were dissected, cleaned, and placed in biopsy cassettes for processing for histology and hematoxylin and eosin (H&E) staining. Histology and H&E slides were prepared by the Texas A and M University Veterinary Medicine and Biomedical Sciences Research Histology Unit Core Facility (RRID:SCR_022201). H&E slides were imaged with a Leica DM 6B upright microscope (Wetzlar, Germany) at the Texas A and M University Microscopy and Imaging Center Core Facility (RRID:SCR_022128).

Cells were also isolated from the tumors using previously established protocols ^27, 31^. Isolated cells were evaluated for LGR5^+^ cancer stem cell expression using flow cytometry following protocols in section 2.4. Cells were also collected for culture until 1 passage was reached whereby flow cytometry of the LGR5^+^ cancer stem cell phenotype was reassessed.

### 2.7 KRAS^G13D^ CRISPR Mutation in RKO Cells

All components for CRISPR procedures were purchased from Integrated DNA Technologies (IDT; Coralville, IA). The HDR donor template and associated Cas9 guide RNA (crRNA) was designed using the Alt-R^TM^ HDR Design Tool with IDT to create a *KRAS^G13D^* mutation^40^. Potential off-target effects of crRNA candidates were analyzed using the Alt-R^TM^ HDR Design Tool^40^, and the crRNA sequences with the fewest off-target sites in the human genome were selected for further analysis. The target sequences of the crRNA and HDR Oligos used in this study are shown in **Supp. Table 1**. Additional components purchased included Alt-R CRISPR-Cas9 tracrRNA, Alt-R S.p. Cas9-GFP V3, Alt-R® Cas9 Electroporation Enhancer, and Alt-R HDR Enhancer V2. Following manufacturer protocols, the guide RNA (gRNA) complex was created by a 1:1 combination of the designed crRNA:tracrRNA. Then, the ribonucleoprotein complex was created with a 1.2:1.0 molar ratio of gRNA:Cas9-GFP supplemented with PBS, per manufacturer’s protocols^40^.

One day prior to electroporation, media was changed in RKO cells under passage number three in cell culture flasks. On the day of electroporation, RKO cells were detached using 0.25% trypsin and 1 million cells were spun down and resuspended in PBS. The ribonucleoprotein complex, electroporation enhancer, and HDR donor oligos (+ and -) were added to the suspended cells. The mixture was transferred to a pre-cooled (4°C) 0.4mm electroporator cuvette (BioRad, Hercules, CA). Electroporation was performed on the BioRad GenePulser XCell with the following protocol parameters: square, 110V, 25ms, 1 pulse. The protocol was run two times and then cells were transferred to the pre-warmed cell culture flask, with culture medium supplemented with HDR Enhancer V2. Cells were cultured in growth media for 48 hours and examined with a Leica DMi8 microscope (Wetzlar, Germany) to verify GFP expression. Cells were allowed to grow for an additional 48 hours prior to experimental use or collection for knock-in validation. To validate the *KRAS^G13D^* mutation, a sample of cells was lysed in RIPA buffer, for subsequent western blot analysis of phosphorylated ERK and total ERK compared to non-electroporated RKO cells (see Section 2.8 for Western Blot procedure).

### 2.8 Western Blot Analysis of ERK Activity

Phosphorylation of ERK was used to determine the activation status of *KRAS*, and to verify CRIPSR mutation methods employed in RKO cells. Cells exposed to static or peristalsis conditions for 24 hours were detached using 0.25% trypsin, collected from PDMS and lysed in 100 µL of Radio-immunoprecipitation assay (RIPA) Buffer supplemented with 1 µL Halt Protease Inhibitor Cocktail. Extracted protein concentration was measured using the Pierce™ BCA Protein Assay Reagent following manufacturer’s protocol for a 96 well format. Following protein quantification, 10 µg of protein from each sample were loaded onto 4-20% gradient polyacrylamide gels (Novex™, ThermoFisher), and separated via electrophoresis. Protein was transferred to a PVDF membrane and blocked with 2.5% bovine serum albumin (BSA). Membranes were probed with primary antibodies (pERK or Total ERK, ThermoFisher) overnight at 4°C, washed with TBST buffer, and probed with an appropriate HRP-conjugated secondary antibody. GAPDH was used as a loading control to determine changes in phosphorylated ERK (pERK) and Total ERK expression among samples. SuperSignal West Pico PLUS Chemiluminescent Substrate (ThermoFisher) was used to visualize bands on a LI-COR C-DiGit Blot Scanner (LI-COR Biosciences, Lincoln, NE). Digital images acquired were processed and densitometry was performed using NIH Image J. Band intensities were normalized by dividing the intensity obtained from each protein band to their corresponding GAPDH band intensity. Band intensities of pERK were normalized with respect to average total ERK band intensity to obtain a ratio of pERK over total ERK.

### 2.9 WNT Pathway Gene Expression Analysis

Following exposure to static or peristalsis conditions for 24 hours, RNA was lysed directly from the PDMS membrane using the RNeasy Mini Kit (Qiagen, Hilden, Germany). RNA concentration and purity were evaluated using a NanoDrop OneC (ThermoFisher Scientific) and stored at −80°C until ready to use. Reverse transcription was performed following manufacturer’s protocols using the High-Capacity cDNA Reverse Transcription Kit (Applied Biosystems). qPCR was performed with a QuantStudio5 (Applied Biosystems) using the Applied Biosystems PowerUp SYBR Green PCR Mastermix (Thermofisher Scientific) for detection. Genes that were investigated included canonical Wnt ligands: *WNT1* (Wnt family member 1), *WNT7b* (Wnt family member 7b), *WNT8a* (Wnt family member 8a), and noncanonical Wnt ligands: *WNT4* (Wnt family member 4), *WNT5a* (Wnt family member 5a), *WNT5b* (Wnt family member 5b). The primer sequences used for each gene are shown in **Supp. Table 2**. Changes in gene expression were calculated using the 2ΔΔCt method, with *GAPDH* as the housekeeping control^41^. qPCR experiments were run in triplicates, with 3-4 independent biological replicates. Comparisons were drawn between static controls and peristalsis.

### 2.10 Immunofluorescent Staining and Fluorometry of β-Catenin

Immunofluorescence of PDMS membranes exposed to peristalsis or maintained as static controls was performed using previously established protocols^25, 26^. Following formalin fixation, and blocking and permeabilization, cell-seeded membranes were incubated with the fluorescently tagged primary antibody βcatenin-AlexaFluor647 and a nuclear counterstain (DAPI) for 1 hour at room temperature. Unbound antibodies were rinsed using PBS and DI water and mounted using the ProLong™ Diamond antifade mounting reagent (ThermoFisher). Fluorescence was observed using an Olympus Fluoview FV3000 Confocal Laser Scanning Microscope (Tokyo, Japan) with 5 independent, non-overlapping regions for analysis. NIH Image J was employed to perform fluorometry. β-catenin activation was quantified as the mean fluorescent intensity of β-catenin (pink) located within the nucleus (blue) (**Supp. Fig. 2**). Mean intensity values from each test condition were compared to their respective static controls to produce a fold change. Microscopy was performed at the Texas A&M Health Science Center Integrated Microscopy and Imaging Laboratory Core Facility (RRID:SCR_021637).

### 2.11 Statistical Analysis

Statistical analysis was performed on GraphPad Prism 10. All reported values result from 3-5 independent biological replicates. In vivo tumor and xenograft harvest analyses result from 10-12 independent biological replicates. All qPCR data was normalized to static conditions within each experimental set and performed in triplicates over at least 3 biological replicates. Image analysis and morphometry included 5 non-overlapping fields of view from 3 biological replicates. Two-way ANOVA, t-test, and one-way ANOVA analyses were performed as appropriate, and statistical significance is indicated within each experimental data set with associated p-values. Box and whiskers plots were used to represent collected data. Box values range from the 25^th^ to 75^th^ percentiles with a line at the median and whiskers extend from the smallest value to the largest value.

## Results

### Peristalsis enhances LGR5 expression in CRC cells harboring activating KRAS mutations

To establish a link between LGR5 expression and *KRAS* activating mutations in CRC, cell and PDX lines were exposed to peristalsis or maintained as static controls for 24 hours prior to flow cytometry analysis of LGR5, a CRC stem cell marker. The gating strategy for all flow analysis is provided in **Supp. Fig. 1**. In HCT116 *KRAS^G13D^*, peristalsis significantly increased LGR5^+^ cells by 3.1-fold compared to static controls (****p<0.0001, two-way ANOVA; **Fig. 1A-B**; representative flow analysis plots and quantification). Similar to HCT116 *KRAS^G13D^*, exposure to peristalsis increased LGR5^+^ expression 2.5-3.8-fold in LS174T *KRAS^G12D^* and PDX1 *KRAS^G12C^* cells (****p<0.0001; two-way ANOVA; **Fig. 1B**). In contrast, exposure of RKO *KRAS^WT^* cells to peristalsis did not increase LGR5 expression compared to static controls (***p<0.001, two-way ANOVA; **Fig. 1C-D**; representative flow analysis plots and quantification). Similarly, *KRAS^WT^* lines, PDX2 (*p<0.05) and PDX3 (ns) resulted in no LGR5 enrichment compared to static controls (two-way ANOVA; **Fig. 1D**).

**Figure 1:**
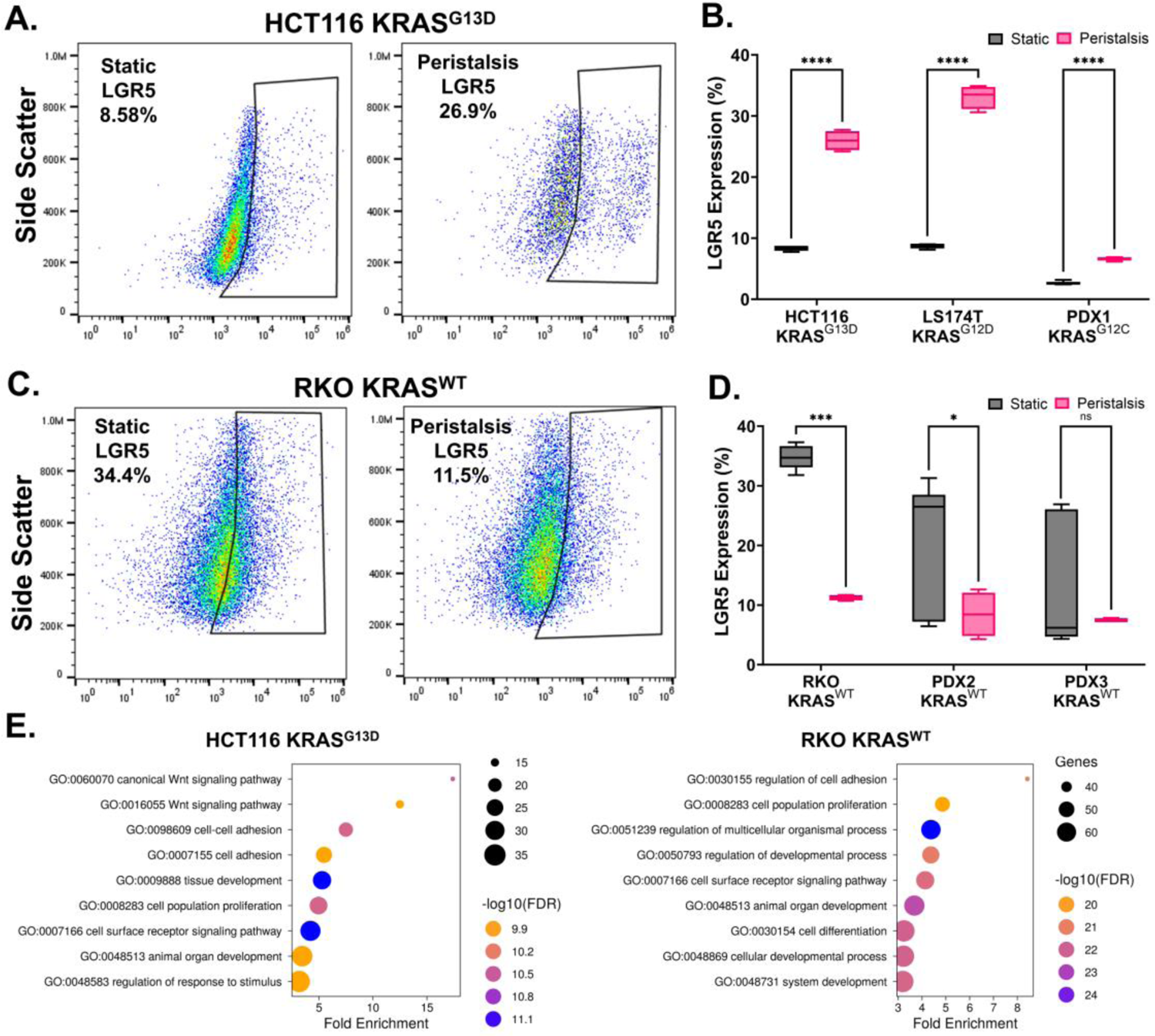
LGR5^+^ cancer stem cell enrichment in *KRAS* mutant CRC cells exposed to peristalsis. (A) Representative LGR5 flow cytometry plots of HCT116 *KRAS^G13D^* cells maintained in static controls or exposed to peristalsis in the bioreactor. (B) Box and whiskers plots summarizing flow analysis of *KRAS* mutant LGR5^+^ expression (%) after 24hr exposure to peristalsis bioreactor or maintenance in static controls. Significant increase in LGR5^+^ expression was noted in all cell types exposed to peristalsis compared to static controls (****p<0.0001, two-way ANOVA). (C) Representative LGR5 flow cytometry plots of RKO *KRAS^WT^* cells maintained in static controls or exposed to peristalsis in the bioreactor. (D) Box and whiskers plots summarizing flow analysis of *KRAS* wild type LGR5^+^ expression (%) after 24hr exposure to peristalsis bioreactor or maintenance in static controls (****p<0.0001, *p<0.05, ns, two-way ANOVA). (E) Gene ontology pathway analysis using Weighted Gene Correlation Network Analysis in HCT116 *KRAS^G13D^* and RKO *KRAS^WT^*cells exposed to peristalsis. The top 9 GO terms from each population are displayed.

To evaluate peristalsis-associated mechanotransduction in HCT116 *KRAS^G13D^* cells compared to RKO *KRAS^WT^*, global RNA-Seq gene expression analysis methods were employed. Gene ontology pathway analysis using Weighted Gene Correlation Network Analysis (WGCNA) between HCT116 *KRAS^G13D^* and RKO *KRAS^WT^* cells identified 6 unique pathways when assessing the top 9 GO terms for each population (**Fig. 1E**). Unbiased bioinformatics analysis demonstrated that HCT116 *KRAS^G13D^* cells showed enrichment in response to stimulus, adhesion, and Wnt signaling gene sets, while RKO *KRAS^WT^* cells showed enrichment for developmental processes, cell differentiation, and proliferation gene sets.

### Peristalsis drives increased tumorigenicity in HCT116 KRAS^G13D^ mutant tumors

We next assessed the role peristalsis plays in driving tumor growth in response to KRAS activating mutation *in vivo*. HCT116 *KRAS^G13D^* and RKO *KRAS^WT^* cells were pre-exposed to peristalsis or static control conditions for 24 hours, then injected subcutaneously into NOD.Cg-Prkdc^scid^ Il2rg^tm1Wjl^/SzJ (NSG) mice (**Fig. 2A**). Representative images of harvested tumors showed overall differences in tumor sizes across conditions (**Fig. 2B**). HCT116 *KRAS^G13D^* tumors initiated with peristalsis-conditioned cells were larger compared to static tumors (**Fig. 2B**). Interestingly, the size differences in peristalsis versus static conditions was not observed in RKO *KRAS^WT^* tumors (**Fig. 2B**).

**Figure 2:**
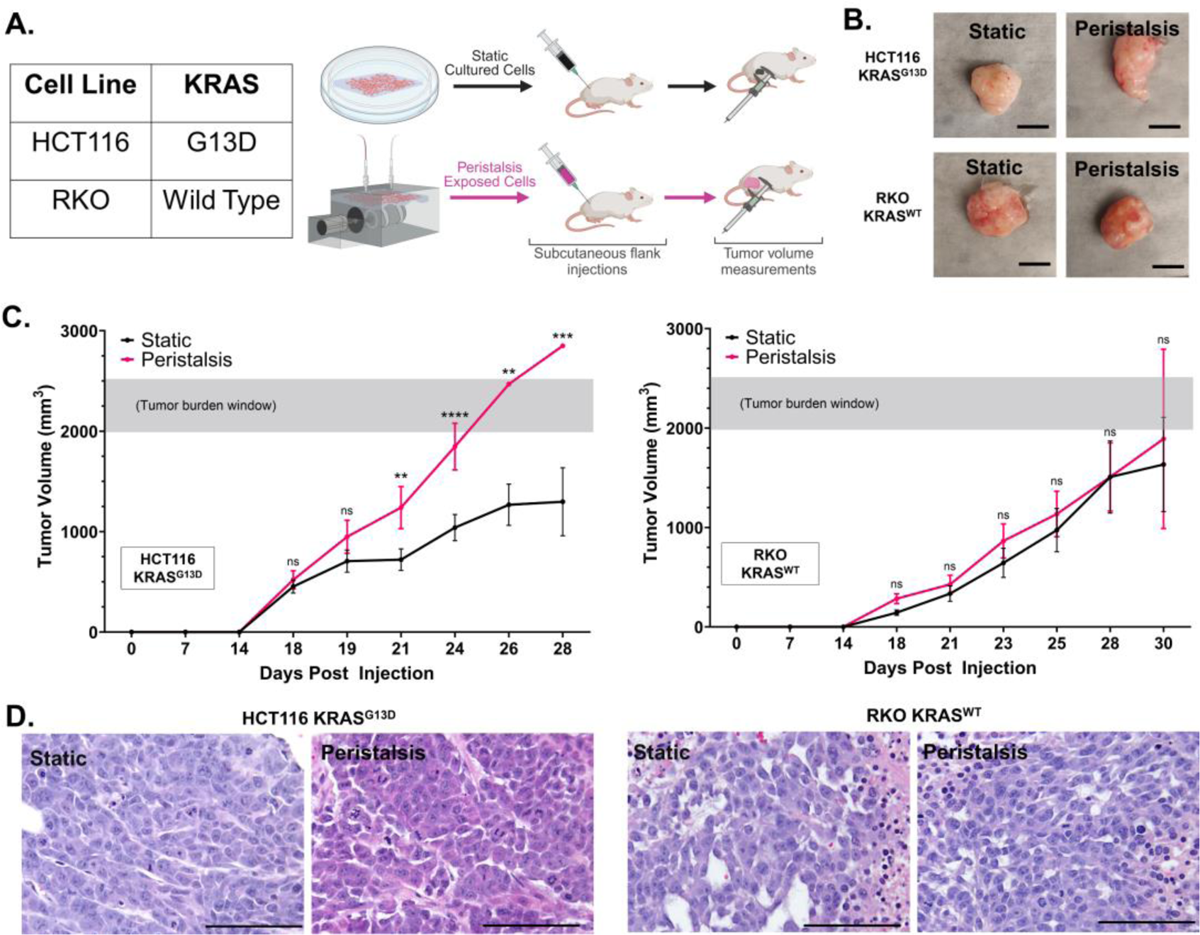
Increased tumor growth in HCT116 *KRAS^G13D^* mutant tumors in peristalsis-conditioned cells. (A) Depiction of in vivo subcutaneous xenograft procedure from cell collection to analysis method. Cell lines used for in vivo studies and their corresponding KRAS status information is included in the table. Cells were cultured statically or exposed to peristalsis for 24 hours prior to collection and subsequent subcutaneous injection into the left and right flank of NSG mice. Mouse weight and tumor volume measurements were continuously monitored. (B) Representative images of harvested tumors from static- and peristalsis-conditioned HCT116 *KRAS^G13D^* and RKO *KRAS^WT^* cells. Scale bar 1cm. (C) Tumor initiation and growth kinetics are shown for both HCT116 *KRAS^G13D^* and RKO *KRAS^WT^*. In HCT116 *KRAS^G13D^*, peristalsis tumors (pink curve) demonstrated elevated tumorigenicity, with faster tumor initiation and tumor burden development (gray shaded area) compared to static tumors (black curve; ns, **p<0.01, ***p<0.001, ****p<0.0001, two-way ANOVA). RKO *KRAS^WT^* tumors demonstrated overlap in growth kinetics with no significant differences in tumor initiation nor development (ns, two-way ANOVA). (D) Histological examination of H&E stains demonstrates that both static and peristalsis tumors in HCT116 *KRAS^G13D^* cells revealed highly invasive cell morphology. Harvested RKO *KRAS^WT^* tumors in peristalsis and static conditions demonstrated less invasive cell morphology with necrosis present. Scale bar 100μm.

To support endpoint visual analysis, growth rates were tracked throughout tumor growth. Overall growth kinetics in the HCT116 *KRAS^G13D^* mice were significantly different between the static and peristalsis conditions (**Fig. 2C**). HCT116 peristalsis tumors developed at a faster rate of 177.1 mm^3^/day while static tumors were 0.58-fold slower at a rate of 103.1 mm^3^/day in the first week of growth (days 14-21). Peristalsis tumors reached the maximum tumor burden window much earlier (Day 24), compared to static tumors in HCT116 (Day 28; **Fig. 2C**). These differences in growth kinetics of the tumors appear as divergent curves between the statis and peristalsis conditions in HCT116 *KRAS^G13D^*.

In the RKO *KRAS^WT^* tumors, the growth kinetics were similar in both the static and peristalsis conditions (almost overlapping static and peristalsis growth curves in **Fig. 2C**). RKO peristalsis tumors grew at a rate of 60.9 mm^3^/day and static tumors were only 0.78-fold slower at 47.8 mm^3^/day in the first week of growth (days 14-21). Both peristalsis and static tumors reached the tumor burden window on Day 30 with little difference in overall growth patterns (**Fig. 2C**). Histologic examination of harvested HCT116 xenografts revealed highly invasive cell morphology in both static and peristalsis conditions (**Fig. 2D**). In harvested RKO xenografts, histological examination demonstrated necrosis and less invasive cell morphology across both static and peristalsis conditions (**Fig. 2D**).

Since there were divergent growth rates in peristalsis conditioned HCT116 *KRAS^G13D^* tumors, we tested the hypothesis that LGR5 enrichment would be seen across the xenograft process. At the termination of the growth studies, harvested xenografts were dissociated and collected for flow LGR5 enrichment analysis on the day of harvest and following the first passage of isolated cells. HCT116 *KRAS^G13D^* xenografts showed increased LGR5 expression in the peristalsis conditioned group, both on the day of harvest and at the time of the first passage (****p<0.0001, two-way ANOVA compared to xenografts from static groups, **Fig. 3A**). LGR5 expression analysis from RKO *KRAS^WT^* xenografts demonstrated no significant differences between static and peristalsis conditions on the day of harvest or at the time of the first passage (ns, two-way ANOVA, **Fig. 3B**). Note that no additional peristaltic mechanical stimulation was provided following xenograft tumor harvests, yet LGR5 enrichment persisted in the HCT116 *KRAS^G13D^* cell line, while it did not in the RKO *KRAS^WT^* cell line.

**Figure 3:**
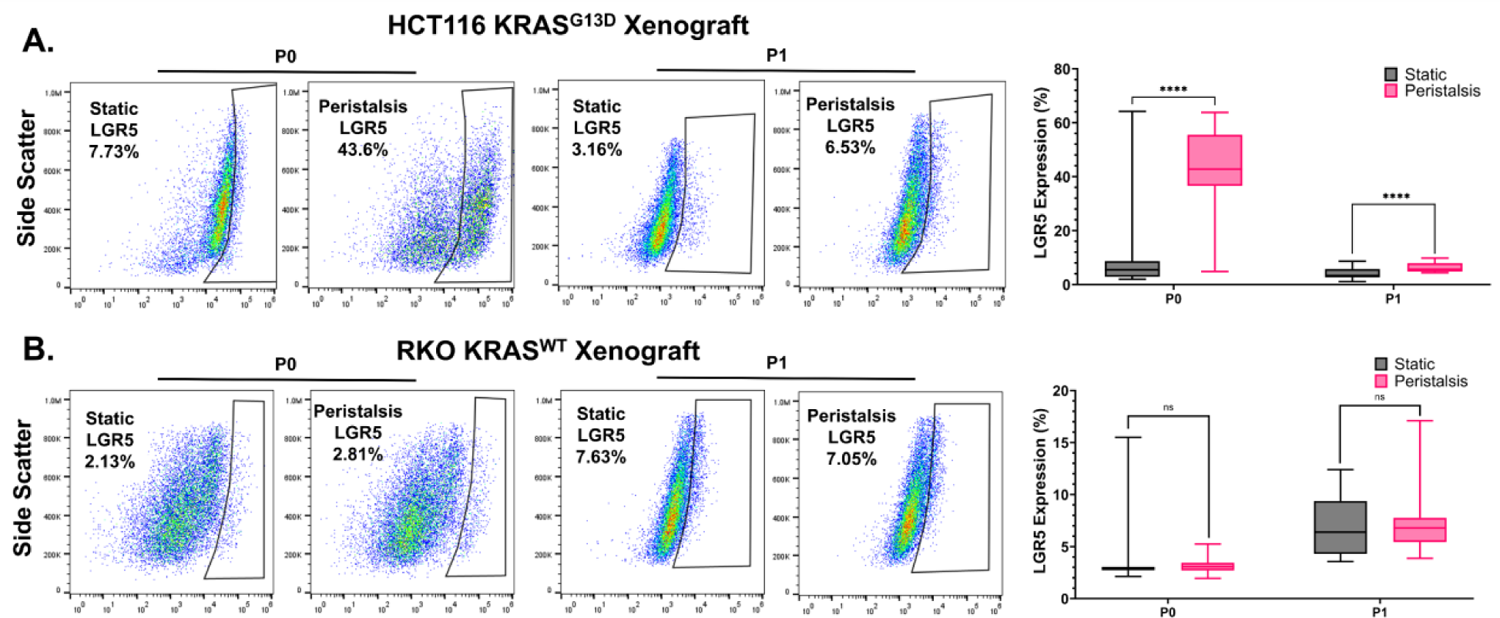
Peristalsis-conditioned HCT116 *KRAS^G13D^* mutant xenografts maintain enriched LGR5^+^ cancer stem cell expression upon harvest and passage. (A) Representative LGR5^+^ flow analysis plots and quantification for harvested HCT116 *KRAS^G13D^* xenografts at passage 0 (P0) and passage 1 (P1). Both P0 and P1 analysis demonstrated increased LGR5 enrichment in peristalsis compared to static-conditioned cells (****p<0.0001, two-way ANOVA). (B) Representative LGR5^+^ flow analysis plots and quantification for harvested RKO *KRAS^WT^* xenografts at passage 0 (P0) and passage 1 (P1). P0 and P1 flow cytometry analysis resulted in no differences between peristalsis and static-conditioned cells (ns, two-way ANOVA).

### The KRAS^G13D^ mutation in WT RKO cells increased LGR5 expression

To test the sufficiency of the G13D *KRAS* mutation in driving LGR5 expression, we generated a CRISPR mediated *KRAS^G13D^* mutation in RKO cells. Pathway activation downstream of the KRAS mutation was functionally validated by western blot analysis of total ERK1/2 versus phosphorylated ERK (pERK1/2). Band intensities were normalized to their corresponding GAPDH loading control. ERK activity was determined as a ratio of pERK over average total ERK. The CRISPR edited RKO *KRAS^G13D^* mutation resulted in a 12% increase in pERK/Total ERK expression compared to RKO *KRAS^WT^* cells (*p<0.05, t-test, **Fig. 4A**). Once increased pERK activity was confirmed as a consequence of introducing the *KRAS^G13D^* mutation into RKO cells, we tested the hypothesis that peristalsis would result in LGR5 enrichment, similar to that observed in the HCT116 *KRAS^G13D^* cell line. As predicted, exposure of mutated RKO *KRAS^G13D^* cells to peristalsis resulted in a 1.73-fold increase in LGR5 expression compared to static controls (***p<0.001, t-test, **Fig. 4B**).

**Figure 4:**
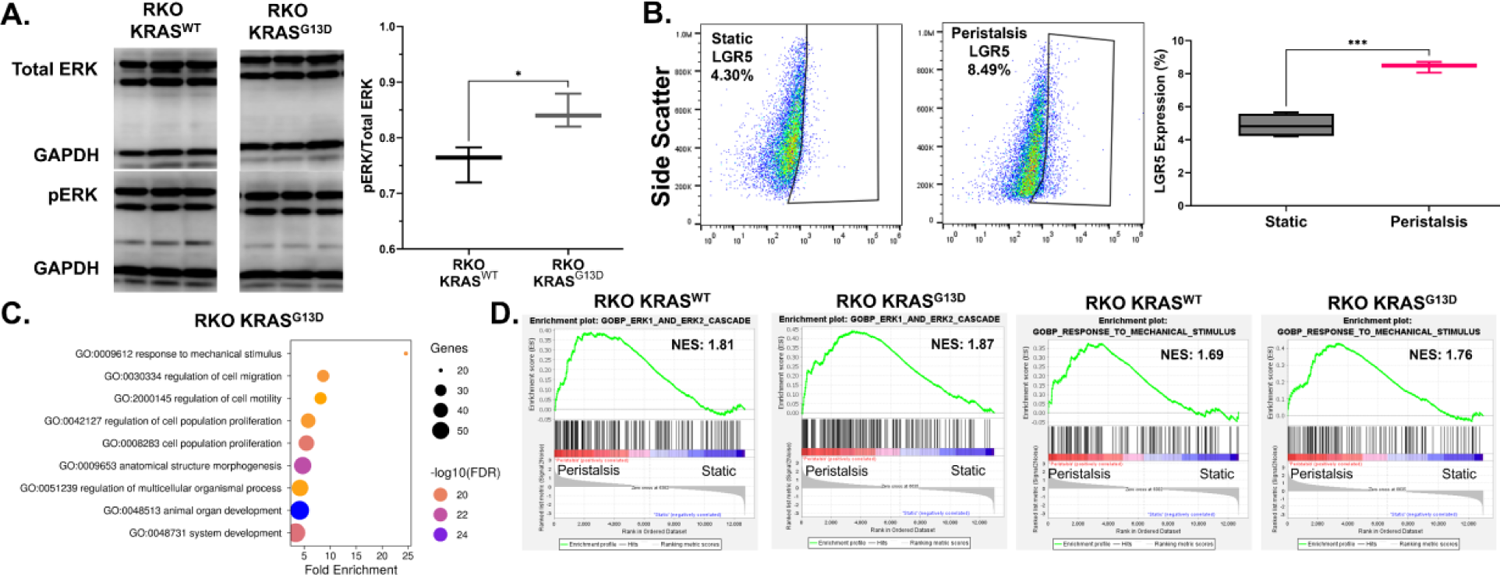
Introduction of the *KRAS^G13D^* mutation in WT RKO cells increases LGR5 expression and alters mechanoresponse. (A) Densiometric analysis of western blots for ERK activity. Band intensities were normalized to their corresponding loading control GAPDH. Phosphorylated ERK (pERK) band intensity was normalized with respect to average total ERK band intensity to obtain a ratio of pERK over total ERK. Introduction of the *KRAS^G13D^* mutation into RKO cells resulted in increased pERK/total ERK expression (*p<0.05, t-test). (B) Representative analysis graphs and box and whiskers plot of flow cytometry LGR5 expression (%) after 24hr exposure to static or peristalsis in RKO *KRAS^G13D^* cells (***p<0.001, t-test). (C) Gene ontology pathway evaluation using Weighted Gene Correlation Network Analysis in RKO *KRAS^G13D^* cells exposed to peristalsis. The top 9 GO terms from the tested population are displayed. (D) Gene set enrichment analysis plots for ERK1_and_ERK2_cascade (GO:0070371) and response_to_mechanical_stimulus (GO:0009612) selected from significantly enriched (FDR < 0.25) gene ontology terms for RKO *KRAS^WT^* and RKO *KRAS^G13D^* cells. Plots include the profile of the running enrichment score and positions of gene set members on the rank-ordered list. Normalized enrichment scores (NES) are also reported where a higher NES indicates more positively enriched genes in the gene set.

In tandem, RNA-seq analysis was employed to evaluate gene expression pathway involvement in peristalsis-associated mechanotransduction in RKO cells with the *KRAS* mutation (**Fig. 4C**). Unbiased analysis with WGCNA plots showed that 5 unique pathways were identified out of the top 9 GO terms between RKO *KRAS^WT^* (**Fig. 1E**) and RKO *KRAS^G13D^* cells (**Fig. 4C**). In the analysis of GO pathways with WGCNA, RKO *KRAS^G13D^* cells showed altered responses from RKO *KRAS^WT^* with significant enrichment in motility, migration, and mechanical response gene sets. Further, Gene Set Enrichment Analysis (GSEA) was performed against the entire gene ontology biological process gene set to look at overall enrichment differences. Closer examination of significant (FDR <0.25), relevant biological processes led to the selection of the ERK1_and_ERK2_cascade (GO:0070371) and response_to_mechanical_stimulus (GO:0009612) plots for comparison (**Fig. 4D**). Interestingly, mutated RKO *KRAS^G13D^* cells had a higher normalized enrichment score (NES) for both gene sets compared to RKO *KRAS^WT^* cells, indicating global changes following the introduction of the *KRAS^G13D^* mutation.

### Peristalsis driven LGR5 enrichment in KRAS^G13D^ cells is independent of Wnt

The Wnt pathway is a known driver of cancer cell proliferation, stemness, and malignant transformation^42^, including driving the enrichment of LGR5. Given the differences in LGR5 expression between *KRAS* WT and G13D lines, we evaluated if there were similar differences in Wnt pathway involvement in *KRAS^WT^* and *KRAS^G13D^* cell populations (**Fig. 5**). We first assessed *Wnt* ligand expression via qPCR gene expression. In the HCT116 *KRAS^G13D^* line, cells exposed to peristalsis significantly increased gene expression of many canonical and noncanonical *Wnt* ligands compared to static controls (*WNT7b*, *WNT4*, *WNT5a*, and *WNT5b*; **Fig. 5A, Supp. Fig. 3**). In contrast, only a mild increase in non-canonical *Wnt* ligands (*WNT4*, *WNT5a*, and *WNT5b*) were observed in RKO *KRAS^WT^* cells as a result of peristalsis, differing quite significantly from the patterns observed in the HCT116 *KRAS^G13D^* cell line (**Fig. 5A, Supp.** Fig. 3). Importantly, when the *KRAS^G13D^* mutation was introduced into RKO, peristalsis again resulted in significant increases in *Wnt* ligand genes similar to the HCT116 *KRAS^G13D^* condition (*WNT1*, *WNT7b*, *WNT4*, *WNT5a*, and *WNT5b*; **Fig. 5A, Supp.** Fig. 3).

**Figure 5:**
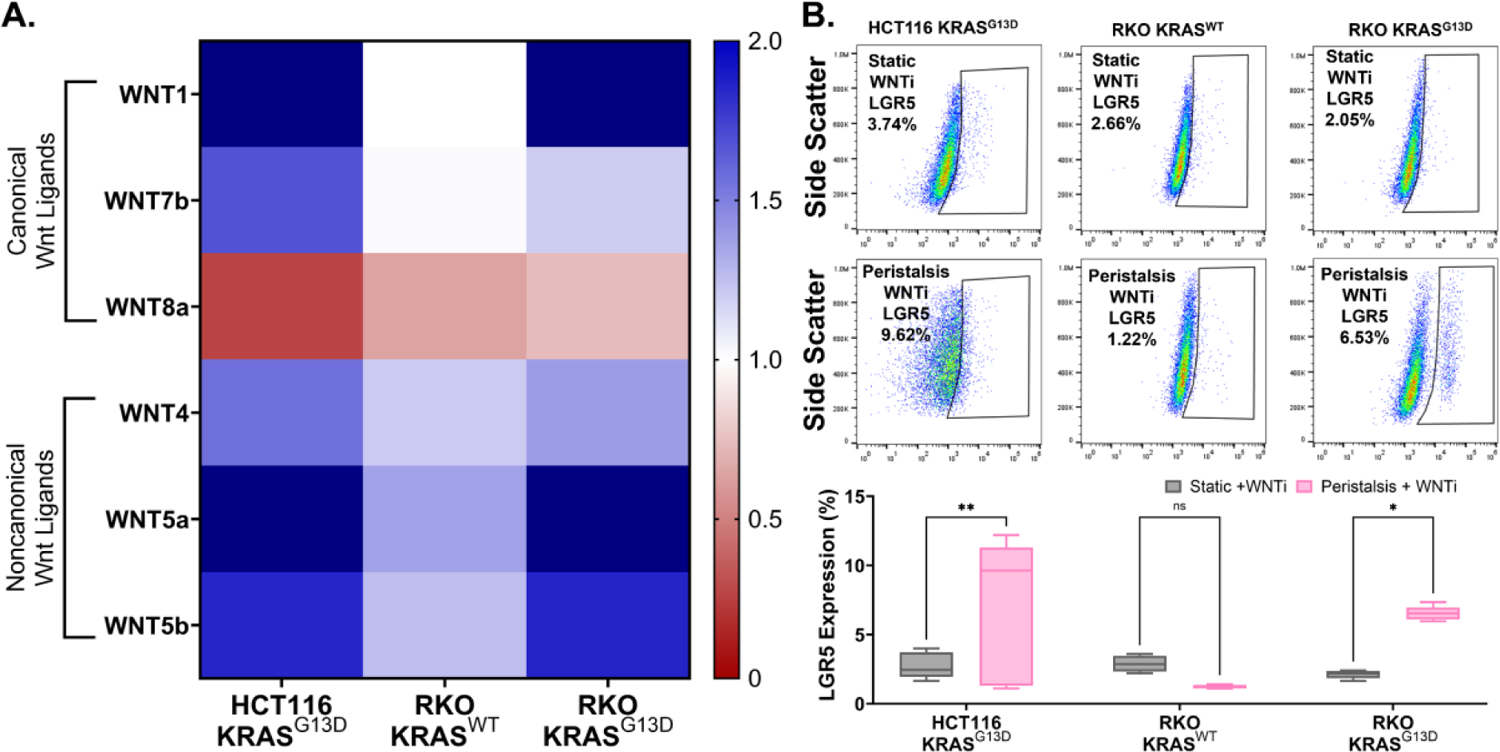
LGR5 enrichment in *KRAS^G13D^* mutant cells in response to peristalsis is independent of Wnt. (A) Heat map of *Wnt* ligand gene expression. Gene expression changes in peristalsis for HCT116 *KRAS^G13D^*, RKO *KRAS^WT^* and RKO *KRAS^G13D^* cells are demonstrated as fold-change values compared to static controls (indicated by 1.0). Statistical differences between each cell type exposed to peristalsis and static controls individually can be found in Supplementary material (**Supp.** Fig. 3). (B) Representative flow cytometry analysis graphs and box and whisker plot of flow cytometry LGR5 expression (%) following 24 hour exposure to peristalsis or static conditions with Wnt inhibition (WNTi). HCT116 *KRAS^G13D^* and RKO *KRAS^G13D^* resulted in sustained increases in LGR5 expression in peristalsis compared to static controls (**p<0.01, *p<0.05, two-way ANOVA).

*Wnt* ligand gene expression trends implicated the involvement of the Wnt pathway in response to peristalsis. This led us to test the hypothesis that inhibiting pan-Wnt secretion with LGK974^43, 44^ would abrogate peristalsis-associated LGR5^+^ enrichment observed in the *KRAS* mutant cell lines. Interestingly, HCT116 *KRAS^G13D^* cells still resulted in a significant 2.5-fold increase of LGR5 expression following peristalsis compared to static controls with Wnt inhibition (**p<0.01, two-way ANOVA, **Fig. 5B**). As expected, neither Wnt inhibition nor peristalsis made a difference in LGR5 levels in RKO *KRAS^WT^* cells (ns, t-test, **Fig. 5B**). Interestingly, despite Wnt inhibition in the *KRAS^G13D^* mutated RKO cells, peristalsis continued to result in a significant 3.1-fold increase in LGR5 expression (*p<0.05, two-way ANOVA, **Fig. 5B**).

### Peristalsis results in Wnt-independent β-catenin activation in KRAS^G13D^ mutant cells

Having demonstrated that LGR5 enrichment downstream of peristalsis was independent of *Wnt* ligand expression in KRAS mutant cells, we then evaluated the activation of the main Wnt pathway effector, β-catenin. We used immunofluorescence to localize β-catenin expression in cells exposed to peristalsis with or without the pan-Wnt secretion inhibitor (WNTi; **Fig. 6**). Activated β-catenin localized to the nuclear region, indicated nuclear translocation of β-catenin (detailed methods for quantification provided in **Supp.** Fig. 2 and representative channel separated images are provided in **Supp.** Fig. 4). All test conditions were normalized to their respective static conditions to report as a fold-change. Exposure to peristalsis significantly increased β-catenin activation by 1.25-fold in HCT116 *KRAS^G13D^* cells, relative to static controls (****p<0.0001, t-test, **Fig. 6A**). RKO *KRAS^WT^* cells resulted in no significant change in peristalsis, relative to static controls (ns, t-test, **Fig. 6A**). Similar to HCT116 *KRAS^G13D^* cells, RKO *KRAS^G13D^* cells demonstrated a 1.60-fold increase in peristalsis exposed cells relative to static controls (****p<0.0001, t-test, **Fig. 6A**). Interestingly, even when Wnt secretion was inhibited, β-catenin activation was sustained by exposure to peristalsis in the mutant HCT116 *KRAS^G13D^* and RKO *KRAS^G13D^* cell lines, suggesting that the β-catenin activation is not being triggered by Wnt (****p<0.0001, t-test, **Fig. 6A**).

**Figure 6:**
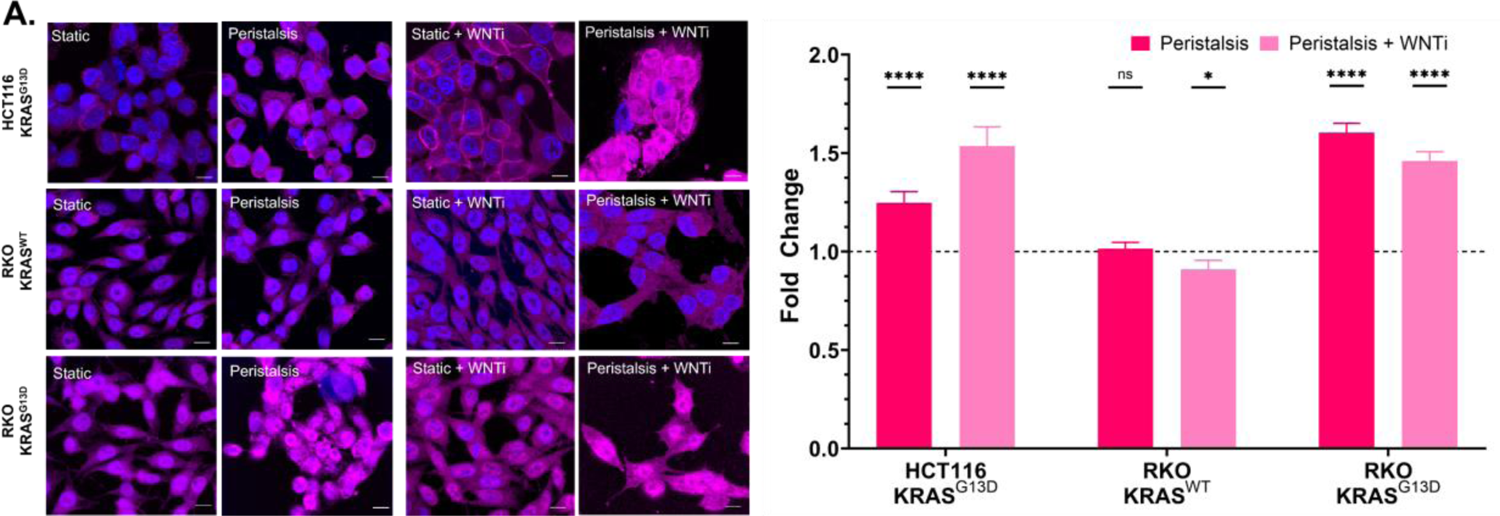
Peristalsis drives Wnt-independent β-catenin activation in *KRAS^G13D^* mutant cells. (A) Representative micrographs of cells maintained as static controls or exposed to peristalsis with and without Wnt inhibition (WNTi) stained with β-catenin (magenta) and counterstained with DAPI (blue). Scale bar 10 µm. Bar graphs quantifying nuclear β-catenin localization relative to their respective static controls for all tested conditions. Peristalsis increased nuclear localization of β-catenin compared to static controls both with and without Wnt inhibition in HCT116 *KRAS^G13D^* and RKO *KRAS^G13D^* (****p<0.0001, t-test). RKO *KRAS^WT^* cells exposed to peristalsis with and without Wnt inhibition did not observe increases in β-catenin nuclear localization. Detailed quantification methods and channel separated images are found in supplementary information (**Supp.** Fig. 2 and 4).

### Peristalsis-associated LGR5 enrichment is sensitive to MEK inhibition

To directly evaluate the dependency of KRAS downstream effectors in regulating LGR5 expression in response to peristalsis, we assessed the effects of Selumetinib (50 nM), a highly selective inhibitor of mitogen-activated protein kinase kinase (MEK) and extracellular signal-regulated kinase (ERK). First, we verified that ERK activity was indeed reduced with selumetinib (evaluating levels of phosphorylated ERK over total ERK). Selumetinib treatment resulted in a 14.4% decrease of phosphorylated ERK activity in HCT116 *KRAS^G13D^* cells compared to uninhibited controls (***p<0.001, t-test, **Fig. 7A**). Although the ERK suppression was modest, we chose this low dose of Selumetinib to preserve cell viability^45, 46^. It was included in the media circulating through the peristalsis bioreactor (MEKi), and cells were collected for analysis of LGR5 expression. Upon MEK inhibition via Selumetinib, exposure to peristalsis no longer resulted in LGR5 enrichment in the HCT116 *KRAS^G13D^* cell line (****p<0.0001, one-way ANOVA, **Fig. 7B**) or RKO *KRAS^G13D^* cells (****p<0.0001, one-way ANOVA, **Fig. 7D**). Additionally, MEK inhibition resulted in dramatically lower LGR5 expression in RKO *KRAS^WT^* cells (****p<0.0001, one-way ANOVA, **Fig. 7C**). Together, these studies link MEK activation to peristalsis-induced enrichment of LGR5.

**Figure 7:**
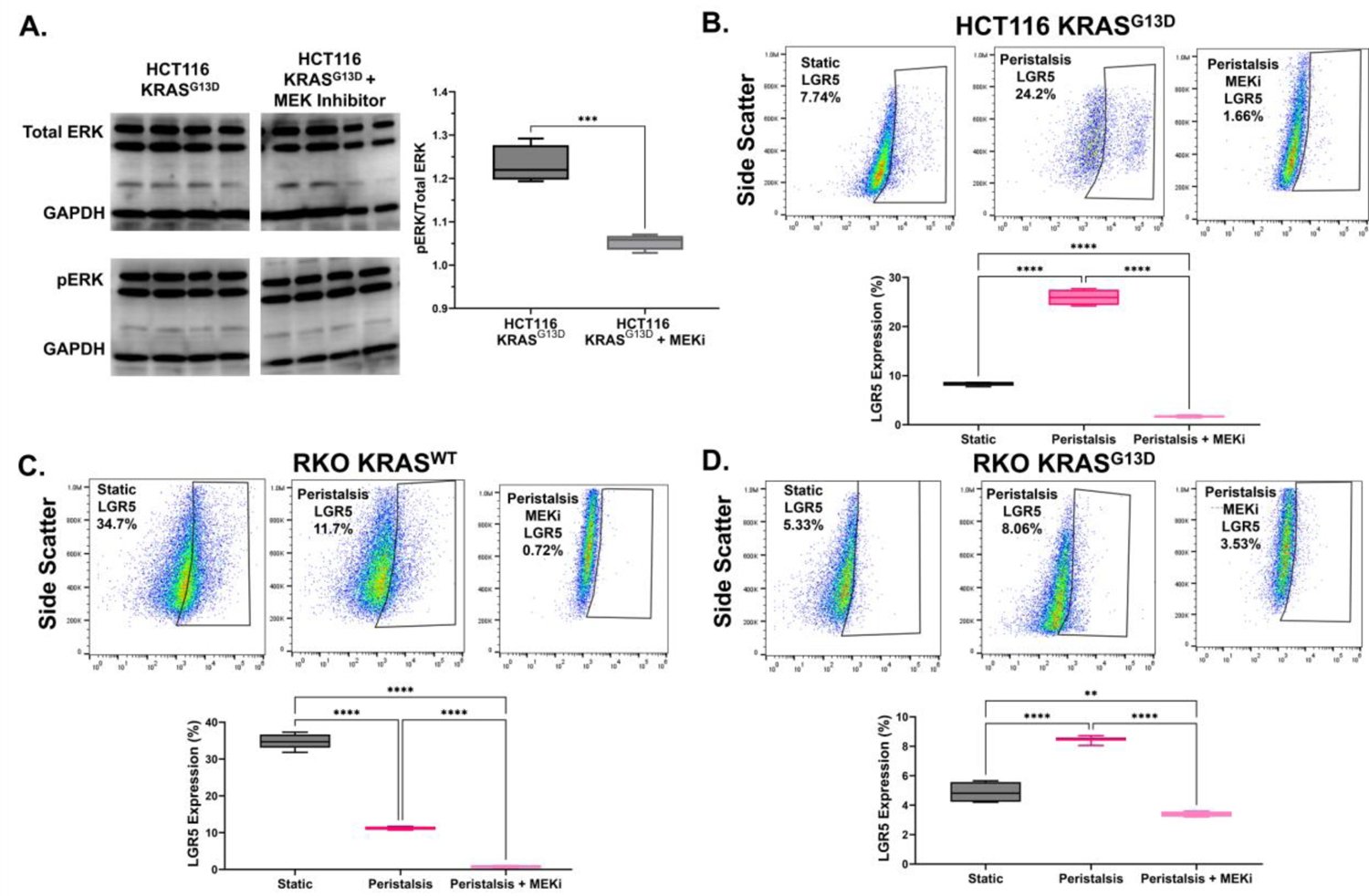
LGR5 enrichment downstream of peristalsis is sensitive to MEK inhibition. (A) Densiometric analysis of western blots for ERK activity analysis in HCT116 *KRAS^G13D^* with and without MEK inhibition (Selumetinib, 50 nM). Band intensities were normalized to their corresponding loading control GAPDH. Phosphorylated ERK (pERK) band intensity was normalized with respect to average total ERK band intensity to obtain a ratio of pERK over total ERK. MEK inhibition in HCT116 *KRAS^G13D^* cells resulted in decreased pERK/total ERK expression (***p<0.001, t-test). Representative flow cytometry analysis graphs and box and whisker plot of flow cytometry LGR5 expression (%) following 24 hour exposure to static, peristalsis or peristalsis with MEK inhibition (MEKi) in (B) HCT116 *KRAS^G13D^*, (C) RKO *KRAS^WT^*, and (D) RKO *KRAS^G13D^* cells. Upon MEK inhibition via Selumetinib, exposure to peristalsis resulted in significantly decreased LGR5 expression for all cell types compared to static or peristalsis alone (**p<0.01, ****p<0.0001, one-way ANOVA).

### MEK inhibition in KRAS^G13D^ mutant cells abrogates peristalsis induced Wnt and β-catenin activation

Since MEK inhibition (MEKi) with Selumetinib abrogated peristalsis-associated LGR5 enrichment in KRAS mutant lines, we explored the effect of MEK inhibition on *Wnt* ligand expression as well as on β-catenin activation. Despite MEK inhibition, peristalsis continued to increase gene expression of many *Wnt* ligands in HCT116 *KRAS^G13D^* cells (*WNT1*, *WNT7b*, *WNT4*, *WNT5a*, and *WNT5b*; **Fig. 8A, Supp.** Fig. 5). Consistent with previous observations, there were no significant *Wnt* ligand increases in the RKO *KRAS^WT^* cells downstream of peristalsis, with MEK inhibition (**Fig. 8A, Supp.** Fig. 5). Introducing the *KRAS^G13D^* mutation into the RKO WT line modestly increased *Wnt* ligand expression downstream of peristalsis with a significant increase in *WNT1* and modest increase in *WNT8a*, *WNT5a*, and *WNT5b* in RKO *KRAS^G13D^*, relative to static controls (**Fig. 8A, Supp.** Fig. 5). Ultimately, MEK inhibition impacted peristalsis induced *Wnt* ligand expression less significantly in *KRAS^G13D^* mutant cells compared to RKO *KRAS^WT^* cells.

**Figure 8:**
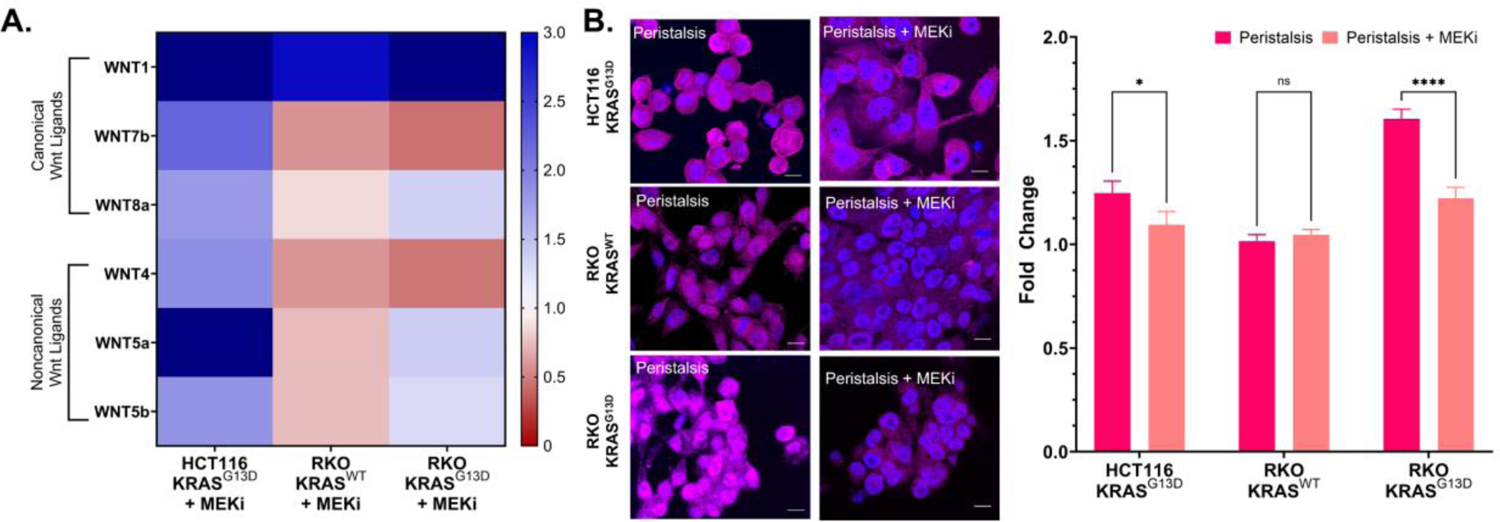
MEK inhibition reduces peristalsis induced Wnt and β-catenin activation in *KRAS^G13D^* mutant cells. (A) Heat map of *Wnt* ligand gene expression. Gene expression changes in peristalsis with MEK inhibition (MEKi; Selumetinib 50 nM) for HCT116 *KRAS^G13D^*, RKO *KRAS^WT^* and RKO *KRAS^G13D^* cells are demonstrated as fold-change values compared to static controls (indicated by 1.0). Statistical differences between each cell type exposed to peristalsis with MEK inhibition and static controls individually can be found in Supplementary material (**Supp.** Fig. 5). (B) Representative micrographs of cells exposed to peristalsis with and without MEK inhibition (MEKi) stained with β-catenin (magenta) and nuclear counterstained with DAPI (blue). Scale bar 10 µm. Bar graphs quantifying nuclear β-catenin localization relative to static untreated controls for all tested conditions. Peristalsis with MEK inhibition decreased nuclear localization compared to peristalsis alone in both HCT116 *KRAS^G13D^* and RKO *KRAS^G13D^* (*p<0.05, ****p<0.0001, two-way ANOVA). RKO *KRAS^WT^* cells exposed to peristalsis with and without MEK inhibition did not observe differences in β-catenin nuclear localization. Detailed quantification methods and channel separated images are found in supplementary information (**Supp.** Fig. 2 and 4).

β-catenin activation in peristalsis was similarly evaluated for its response to MEK inhibition (**Fig. 8B, Supp.** Fig. 4). Compared to peristalsis alone, MEK inhibition resulted in a 12.3% decrease of β-catenin activation in *KRAS^G13D^* HCT116 cells (*p<0.05, two-way ANOVA, **Fig. 8B**). RKO *KRAS^G13D^* cells pheno-copied this sensitivity to MEK inhibition, with a 23.9% decrease in β-catenin activation observed during uninhibited peristalsis (****p<0.0001, two-way ANOVA, **Fig. 8B**). Our data strongly suggest the involvement of the activating *KRAS* mutation in sustaining β-catenin activation downstream of peristalsis, since RKO *KRAS^WT^* cells displayed no difference with or without MEK inhibition.

## Discussion

Mechanics of the colorectal tumor microenvironment (peristalsis) are complex, with propagating waves of multi-axial strain and shear forces always present^23, 24^. In order to study mechanotransduction in response to colonic peristalsis, we previously custom-built a peristalsis bioreactor that mimicked mechanical patterns of the colon *in vitro*^25^. Leveraging the peristalsis bioreactor, we demonstrated that peristalsis resulted in malignant progression of colorectal cancer cells, enriching cancer stem cells and increasing invasive features in mechanically stimulated cells^26^. Interestingly, the mechanotransductive response to peristalsis was specific to cancer cells that carried activating *KRAS* mutations, since LGR5^+^ enrichment was not observed in mechanically stimulated non-cancerous intestinal epithelial cells nor *KRAS* wild type cancer cells. Overall, this led us to hypothesize that activating *KRAS* mutations might alter mechano-responsiveness of colorectal cancer cells to peristalsis.

Oncogenic *KRAS* mutations (G13D, G12D, G12C, etc.) lead to malignant transformation via the constant activation of the mitogen-activated protein kinase (MAPK) cascade^14, 15^. Activation of RAS (through oncogenic mutations or extracellular signals) results in sequential activation of Raf and MEK kinases, ultimately leading to the phosphorylation and activation of extracellular signal-regulated kinase (ERK)^47^. In CRC, oncogenic KRAS mutations are associated with decreased patient survival^18–20^. Interestingly, in breast and pancreatic cancers, oncogenic *KRAS* alters cancer cell mechanosensing^21^. Given the near constant exposure of colorectal cancer cells to colonic peristalsis, we hypothesized that a similar *KRAS*-specific mechanosensitive phenotype might exist in CRC.

First, we focused on Leucine-rich repeat-containing G-protein-coupled receptor 5 (LGR5), due to previous experiences of having observed that peristalsis resulted in LGR5 enrichment in colorectal cancer cells^26^. Indeed, our data demonstrated that LGR5 enrichment downstream of peristalsis was not universal and confined to oncogenic *KRAS* mutant cells (**Fig. 1**). In fact, when mutant *KRAS^G13D^* was induced into a WT background cell, exposure to peristalsis led to LGR5 enrichment (in RKO *KRAS^WT^* vs. RKO *KRAS^G13D^* cells; **Fig. 1, 4**). This data was also backed by observations from unbiased RNA-Seq analysis, where there were 6 unique pathways between HCT116 *KRAS^G13D^* and RKO *KRAS^WT^* and 5 unique pathways between RKO *KRAS^WT^* and RKO *KRAS^G13D^* out of the top 9 GO terms in response to peristalsis. Our results strongly suggest that oncogenic *KRAS* altered mechanosensing in response to peristalsis. These results are in line with evidence that oncogenic RAS signaling alters cytoskeletal organization and actomyosin contractility in epithelial cells^48^. Both these avenues play important roles in cellular mechanotransduction, implicated in cellular ability to sense strain^49^. In fact, emerging evidence suggests that oncogenic RAS increases the elastic modulus of many types of cancer cells, conferring growth advantages in confined spaces via altered mechanosensing^50, 51^. To the best of our knowledge, our work is the first instance where we report that oncogenic RAS signaling alters mechanotransductive signaling in response to a tissue-level mechanical stimulus, like colonic peristalsis.

Having established the role of oncogenic KRAS-specific mechanotransduction leading to LGR5 enrichment, we then evaluated if a functional advantage was also present. Xenografts generated from peristalsis-exposed HCT116 *KRAS^G13D^* cells grew at a significantly faster rate compared to RKO *KRAS^WT^* cells, likely due to the significant LGR5 enrichment due to peristalsis (**Fig. 2, 3**). This was unsurprising, since the LGR5^+^ population is associated with increased tumor growth, invasion, and therapy resistance in CRC clinical samples^52–55^.

In many studies of oncogenic KRAS related mechanosensing, the focus has typically stayed on cell response to environmental stiffness^50^. Since our readout of malignant progression was focused on quantifying increases in LGR5 expression, we focused on the intersection of the Wnt pathway and oncogenic KRAS. Our motivations were two-fold: i) our previous work demonstrated a connection between Wnt activation and subsequent LGR5 expression^26^; and ii) LGR5 is a target gene of Wnt signaling^56–58^. As expected, *Wnt* ligand gene expression was highly increased in response to peristalsis in HCT116 *KRAS^G13D^* (**Fig. 5A**). These results were somewhat unsurprising as HCT116 cells harbor a *CTNNB1* mutation that prevents the degradation of β-catenin leading to enhanced Wnt pathway signaling^59–61^. While RKO *KRAS^WT^* cells demonstrated mild changes in *Wnt* ligands, the introduction of the *KRAS^G13D^* into RKO cells (despite being *CTNNB1^WT^*) resulted in a similar genotype to HCT116 cells in response to peristalsis. Wnt/β-catenin signaling is activated in response to mechanics in many non-epithelial cell types^62^. For example, mechanical stretch in skeletal osteoblasts and oscillatory shear stress in lymphatic endothelial cells have been cited to increase Wnt/β-catenin signaling and their downstream targets^63, 64^. Our results demonstrate that *Wnt* ligand increases occur downstream of peristalsis mechanics in epithelial cancer cells, but only in those that harbor oncogenic *KRAS* mutations. Surprisingly, inhibiting all Wnt ligand secretion via a porcupine-inhibitor^43, 44^ was insufficient in altering LGR5 enrichment in response to peristalsis (**Fig. 5B**).

Importantly, peristalsis resulted in β-catenin nuclear translocation independent of Wnt inhibition, but only in cells that harbored oncogenic *KRAS* mutations (**Fig. 6**). This phenomenon was independently tested with both a gain of function mutation in the RKO *KRAS^WT^* cell line, and with inhibition of MEK/ERK in the HCT116 *KRAS^G13D^*, RKO *KRAS^WT^*, and RKO *KRAS^G13D^* cell lines. Our observations are in line with reported evidence that cytoskeletal stiffness-mediated mechanics can trigger mechanical activation of β-catenin in cancer cells independent of Wnt ligands^62^. This Wnt-independent mechanical activation of β-catenin subsequently led to cancer stem cell enrichment. Similarly, another report demonstrated that external solid mechanical stresses from the growing colorectal tumor microenvironment also activate β-catenin, leading to enhanced tumor growth^65^. Our data corroborates that β-catenin activation occurs in response to the mechanics of peristalsis, but only in cells harboring oncogenic *KRAS* mutations. The intersection of oncogenic RAS and β-catenin activation has also previously been reported in many different cancers either via PI3K/AKT or MEK/ERK activation^66–70^. Hyper-activation of ERK1/2 or PI3K/AKT observed due to oncogenic KRAS requires β-catenin stabilization and activation in melanoma^67^. Interestingly, RAS-driven hyper-activation and malignant progression in thyroid cancers depend on Wnt-independent β-catenin activation^70, 71^. These reports reinforce our findings that peristalsis-driven, Wnt-independent β-catenin activation is unique to cells that harbor activating oncogenic *KRAS* mutations, where we see hyperactivation of ERK.

Lastly, the LGR5 enrichment and β-catenin activation phenotype downstream of peristalsis was sensitive to MEK inhibition in *KRAS* mutant cells (**Fig. 7-8**). While this confirms the role of oncogenic KRAS itself, it also opens out the possibility of MEK/ERK as potential druggable mechano-sensors in colorectal cancer. Evidence of their mechanosensory activity is found in non-epithelial cell literature, where cardiomyocytes and chondrocytes independently exposed to strain phosphorylate MEK/ERK^72–74^. To the best of our knowledge, this is the first study that links MEK to a mechano-sensory response in KRAS mutant colorectal cancer, especially in response to the native mechanics of colonic peristalsis.

## Conclusion

Collectively, this work supports the hypothesis that oncogenic *KRAS* mutations dictate a unique mechanotransduction response to peristalsis in colorectal cancer (CRC). With the application of peristalsis forces, CRC cells harboring a *KRAS* mutation demonstrated an altered functional phenotype and genotype that portends to increased tumor growth *in vivo*. Importantly, *KRAS* targeted therapeutic options are growing in the treatment of *KRAS* mutant cancers and can be used uniquely with other pharmacologic modulators that regulate colonic peristalsis to achieve favorable therapeutic outcomes. Our work supports the idea that the combination of peristalsis modulation and targeted *KRAS* therapeutics can be employed to modulate CRC progression and thereby increase patient survival.

## Data Availability Statement

Data supporting the findings of this study are deposited into the Texas Data Repository. RNA-Seq data will be deposited to Gene Expression Omnibus under “Oncogenic *KRAS* Mutations Confer a Unique Mechanotransduction Response to Peristalsis in Colorectal Cancer Cells”.

## Acknowledgements and Funding Statement

This work was supported by the Cancer Prevention and Research Institute of Texas (CPRIT) grant RP230204 (SAR) through the Regional Excellence Center in Cancer program at Texas A&M University. This work was additionally supported by NCI R37CA26922 (SAR) and NIGMS R35GM137976 (ANS). The authors acknowledge the support of Dr. Yava Jones-Hall from Texas A&M University for assistance with xenograft histological assessment (Texas A&M University Veterinary Medicine and Biomedical Sciences Research Histology Unit Core Facility; RRID:SCR_022201) and Dr. Preeti Kanikarla at MD Anderson for generating the PDX lines used in this study. Authors also acknowledge the assistance of Arpita Mohaptra for RNA-Sequencing sample quality and quantity analysis and the use of the Texas A&M Molecular Genomics Core for analysis instruments. We acknowledge our use of the gene set enrichment analysis, GSEA software, and Molecular Signature Database (MSigDB) (http://www.broad.mit.edu/gsea/)^38^.

