## Supplementary information for "Oncogenic *KRAS* Mutations Confer a Unique Mechanotransduction Response to Peristalsis in Colorectal Cancer Cells"

**Supplementary Material**

*Flow Cytometry Gating Technique*

Flow cytometry analysis was completed with FlowJo (Ashland, OR). Polygon gates were drawn on isotype flow cytometry graphs to establish a background gate at 0.5%. The same gate was applied to antibody-stained cells to establish a percent positive population.

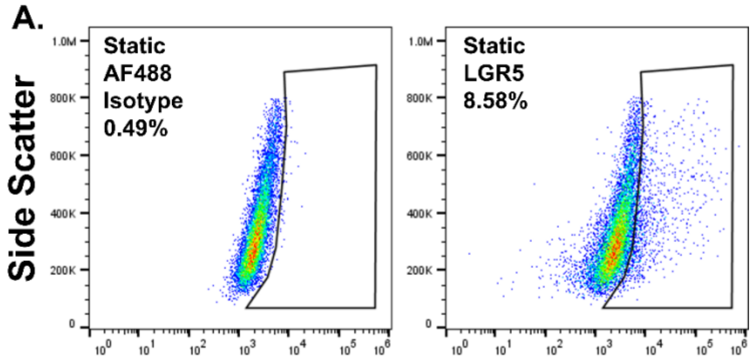


**Supplementary Figure 1: LGR5 flow cytometry gating technique using isotype controls.** A) Polygon gate (outlined in black) was drawn on FlowJo to create a 0.5% background cut off gate in the isotype control graph (left). The same gate was applied to the antibody-stained graph (right) to establish the percentage of the population expressing LGR5.

*CRISPR Reagents*

**Supplementary Table 1:** Target Sequences of CRISPR Reagents created using the Alt-R$^{TM}$ HDR Design Tool.

| CRISPR Reagent | Sequence |
|---|---|
| crRNA | /AlTR1/rCrUrGrArArUrUrArGrCrUrGrUrArUrCrGrUrCrArGr UrUrUrUrArGrArGrCrUrArUrGrCrU/AlTR2/ |
| Alt-R HDR Donor Oligo + | /AlT-R-HDR1/T*C*ATATTCGTCCACAAAATGATTCTGAATTAGC TGTATCGTCAAGGCACTCTTGCCTACGCCATCAGCTCCAACT ACCACAAGTTTATATTCAGTCATTTTCA*G*C/AlT-R-HDR2/ |
| Alt-R HDR Donor Oligo - | /AlT-R-HDR1/G*C*TGAAAATGACTGAATATAAACTTGTGGTA GTTGGAGCTGATGGCGTAGGCAAGAGTGCCTTGACGATAC AGCTAATTCAGAATCATTTTGTGGACGAATAT*G*A/AlT-R-HDR2/ |

*qPCR Wnt Ligand Gene Sequences*

**Supplementary Table 2:** List of genes and corresponding forward and reverse primer sequences used in qPCR amplification.

| Gene | Forward Sequence | Reverse Sequence |
|---|---|---|
| GAPDH | CTGGGCTACACTGAGCACC | AAGTGGTCGTTGAGGGCAATG |
| WNT1 | CGATGGTGGGGTATTGTGAAC | CCGGATTTTGGCGTATCAGAC |
| WNT7b | GAAGCAGGGCTACTACAACCA | CGGCCTCATTGTTATGCAGGT |
| WNT8a | GAACTGCCCTGAAAATGCTCT | TCGAAGTCACCCATGCTACAG |
| WNT4 | AGGAGGAGACGTGCGAGAAA | CGAGTCCATGACTTCCAGGT |

| | | |
|---|---|---|
| WNT5a | TCGACTATGGCTACCGCTTTG | CACTCTCGTAGGAGCCCTTG |
| WNT5b | CATGGCCTACATAGGGGAGG | CTGTGCTGCAATTCCACCG |

*Manual Quantification of β-Catenin Nuclear Localization*

Following Section 2.10, β-Catenin nuclear localization (i.e. activation) was quantified as the mean fluorescent intensity of β-Catenin (magenta) located within the nucleus (blue). Using ImageJ, a mask was created by outlining the nuclei on the DAPI separated channel (outlined with yellow lines for ease of visualization, **Supp. Fig. 2A**). The mask was applied to the β-Catenin separated channel and the “Measure” (CRTL+M) analysis tool was selected to produce a mean intensity value (outlined in Red, **Supp. Fig. 2B**). Mean intensity values from peristalsis conditions were normalized to produce a fold change relative to static control values.

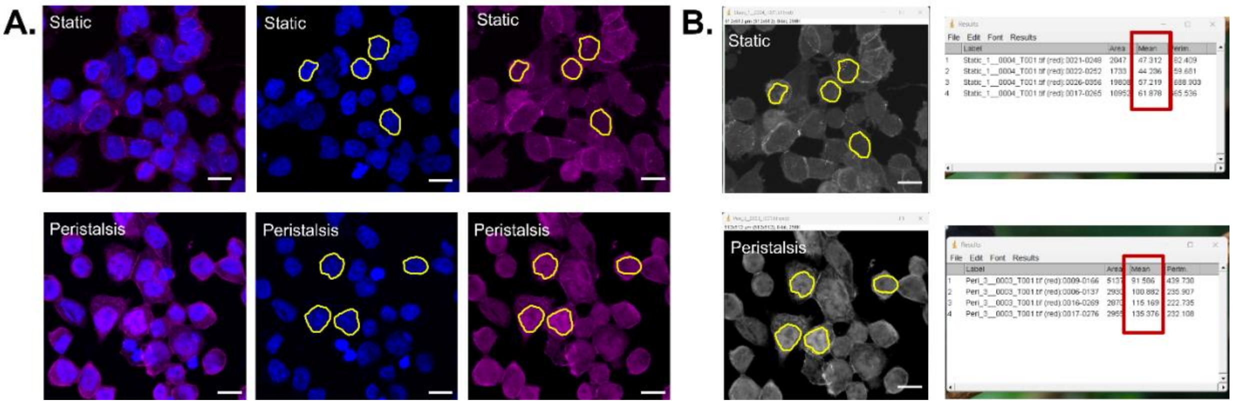


**Supplementary Figure 2: Representative images of manual quantification technique of β-Catenin.** A) Representative micrographs of β-Catenin (magenta) nuclear counterstained with DAPI (blue) following exposure to peristalsis or maintenance of static controls. Yellow outline indicates nuclear mask created using ImageJ. Scale bar 10 µm. B) Screen capture of ImageJ output results used for quantifying mean intensity of β-Catenin inside the nucleus.

*Gene Expression Graphs of Wnt Pathway Genes in Peristalsis*

Breaking down the heat map in **Figure 5**, statistical differences of peristalsis in all three cell types are seen in **Supp. Fig. 3**. Briefly, peristalsis increased expression compared to static controls in *KRAS*$^{G13D}$ HCT116 cells in *WNT7b*, *WNT4*, *WNT5a*, and *WNT5b* (****p<0.0001, t-test). Further, *KRAS*$^{WT}$ RKO cells demonstrated significant increases in *WNT5a* and *WNT5b* (**p<0.01) and *WNT4* (***p<0.001) while *KRAS*$^{G13D}$ RKO cells resulted in significant increases in *WNT1*, WNT7b, *WNT4*, *WNT5a*, and *WNT5b* (*p<0.05, t-test).

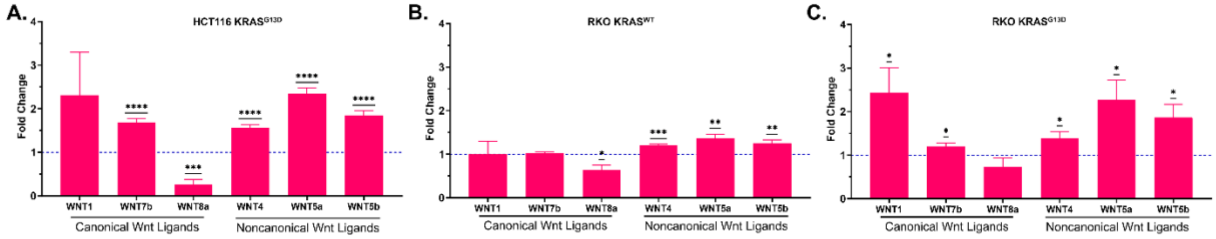

**Supplementary Figure 3: Gene expression analysis of *Wnt* ligand genes in response to peristalsis relative to static controls for *KRAS$^{G13D}$* HCT116, *KRAS$^{WT}$* RKO, and *KRAS$^{G13D}$* RKO cell types.** Bar graph of *Wnt* ligand gene expression analysis in response to peristalsis in A) *KRAS$^{G13D}$* HCT116, B) *KRAS$^{WT}$* RKO, and C) *KRAS$^{G13D}$* RKO cells. Static controls are represented by the blue dotted line at 1, with changes indicated as a relative fold increase compared to static control (*p<0.05, **p<0.01, ***p<0.001, ****p<0.0001, t-test).

*Channel Separated Images of B-Catenin Immunofluorescence*

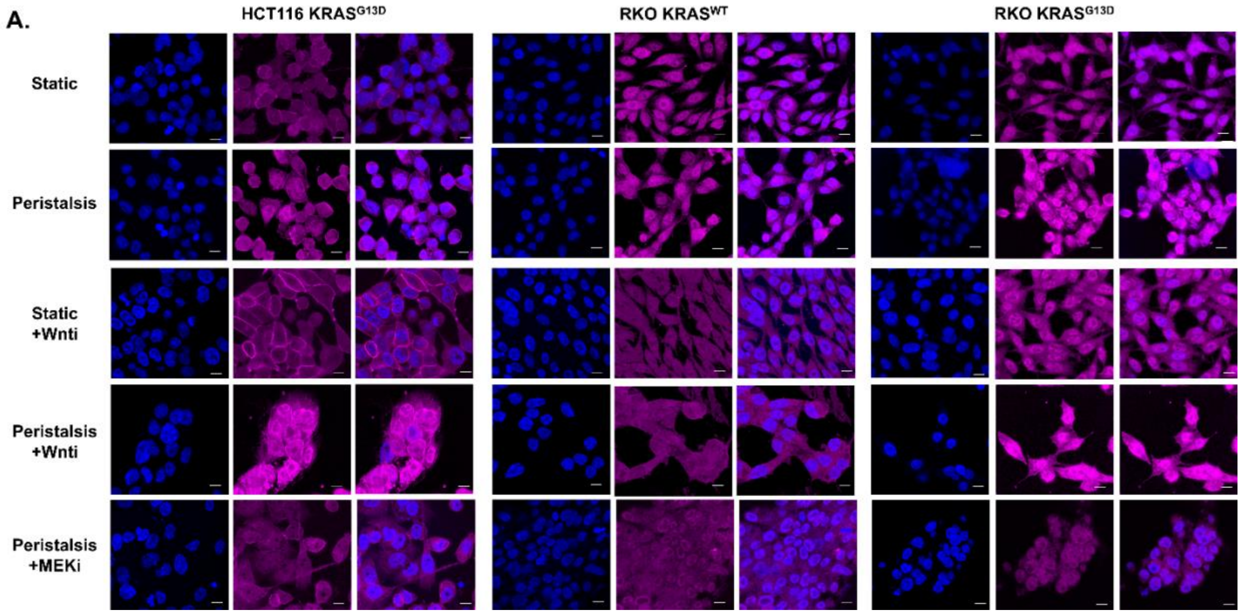


**Supplementary Figure 4: Channel separated images of representative β-Catenin micrographs.** A) Representative micrographs of *KRAS$^{G13D}$* HCT116, *KRAS$^{WT}$* RKO, and *KRAS$^{G13D}$* RKO cells exposed to static or peristalsis conditions with and without inhibitors. Cells stained with β-Catenin (magenta) and nuclear counterstained with DAPI (blue). Scale bar 10 µm.

*Gene Expression Graphs of Wnt Pathway Genes in Peristalsis with MEK Inhibition*

Breaking down the heat map in **Figure 8**, statistical differences of peristalsis in all three cell types are seen in **Supp. Fig. 5**. Briefly, peristalsis with MEK inhibtion increased expression compared to static controls in *KRAS$^{G13D}$* HCT116 cells in *WNT1* (*p<0.05), *WNT5a* and *WNT5b* (**p<0.01) and *WNT7b* and *WNT4* (****p<0.0001, t-test). Further, *KRAS$^{WT}$* RKO cells demonstrated no significant increases in *Wnt* ligand genes while *KRAS$^{G13D}$* RKO cells resulted in a significant increase in *WNT1* (*p<0.05, t-test).

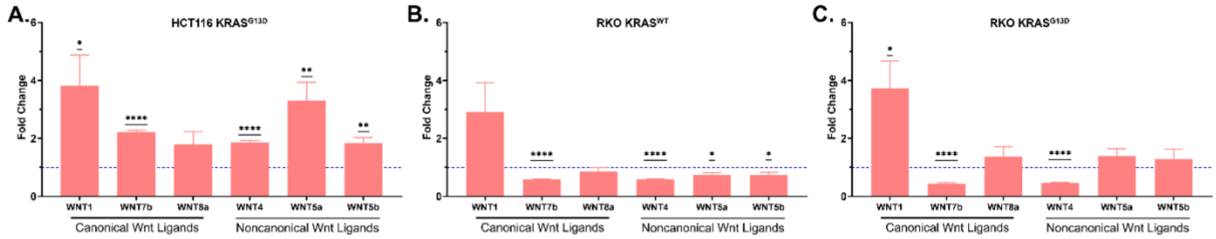

**Supplementary Figure 5: Gene expression analysis of *Wnt* ligand genes in response to peristalsis with MEK inhibition relative to static controls for *KRAS*$^{G13D}$ HCT116, *KRAS*$^{WT}$ RKO, and *KRAS*$^{G13D}$ RKO cell types.** Bar graph of *Wnt* ligand gene expression analysis in response to peristalsis with MEK inhibition in A) *KRAS*$^{G13D}$ HCT116, B) *KRAS*$^{WT}$ RKO, and C) *KRAS*$^{G13D}$ RKO cells. Static controls are represented by the blue dotted line at 1, with changes indicated as a relative fold increase compared to static control (*p<0.05, **p<0.01, ****p<0.0001, t-test).
